# The functional significance of EEG phase synchronization networks during information integration of left and right visual fields

**DOI:** 10.64898/2026.08.21.746382

**Authors:** Makoto Haghihara, Kazumasa Uehara, Yuka O Okazaki, Keiichi Kitajo

## Abstract

Objects moving between the left and right visual hemifields are naturally perceived as continuous entities, although early visual processing independently transmits information from the two hemifields. Therefore, interhemispheric integration of visual information is essential for maintaining an object’s identity. Additionally, brain function is thought to be maintained through a dynamic balance between integration and segregation. In this study, we investigated the functional neural architecture underlying visual hemifield integration in healthy adults, using electroencephalography (EEG) and a visual integration task. To capture neural oscillatory networks without relying on prior assumptions regarding electrode pairs or frequency bands, we applied a frequency-inclusive, data-driven network analysis based on an extended network-based statistic. This analysis identified a broadband EEG phase synchronization network that emerged specifically under task conditions with high interhemispheric integration demands. Furthermore, individual differences in behavioral performance were associated with modulation of interhemispheric synchronization, with this relationship differing according to participants’ relative performance across task conditions. These findings suggest that visual hemifield integration is supported by large-scale phase synchronization networks spanning multiple frequencies and are consistent with the importance of a balance between integration and segregation.

Author Summary

The brain continuously integrates visual information across the left and right visual fields, although early visual processing processes these inputs separately. Efficient brain function is also thought to depend on the balance between the integration and segregation of neural activity. However, the mechanism by which large-scale brain networks achieve this balance during visual information processing remains unclear. In this study, we used EEG and a visual tracking task to examine the neural dynamics during visual hemifield integration. Data-driven network analysis revealed a broadband phase synchronization network that emerged when the interhemispheric integration demands were high. Importantly, individual differences in task performance were associated with the strength of the interhemispheric synchronization. These results suggest that flexible changes in large-scale neural networks across multiple frequencies support visual information integration.

## Introduction

In neural systems, phase synchronization of neural activity has been proposed as a mechanism that enables communication between distant brain regions [1]. Previous studies have demonstrated that the degree of neural phase synchrony increases during a wide range of cognitive tasks [2,3], and is now regarded as a key mechanism through which multiple brain regions form task-relevant functional networks [4,5]. In the context of visual hemifield integration, when participants track objects that move across the left and right visual fields, the visual inputs from the left and right hemifields are independently transmitted to the contralateral primary visual cortices [6,7], making interhemispheric information integration necessary. Indeed, increased interhemispheric EEG phase synchronization—particularly between homologous regions—has been observed when participants track objects moving across the left and right visual fields [8,9]. These findings suggest that EEG phase synchronization facilitates the integration of distributed visual information by enabling coordinated neural communication between distant regions.

Furthermore, it is increasingly recognized that cortico-cortical communication supporting higher-order functions such as integration has been suggested to involve parallel contributions of EEG phase synchronization across multiple frequency bands [10,11]. The hypothesis that these distinct frequencies operate in a hierarchically coordinated manner has gained theoretical support, suggesting that large-scale synchronization networks are organized across multiple frequencies [4]. However, such spatially and spectrally distributed patterns of phase synchronization are difficult to detect using traditional analyses that rely on predefined electrode pairs, specific frequency bands, or homologous region comparisons alone [12].

Although previous studies have primarily focused on the aspect of integration, recent studies have increasingly highlighted segregation—that is, the decoupling or differentiation of processing streams—as an equally important neural mechanism that complements integration in higher-order brain functions [13–16]. In this view, the brain achieves flexible information processing by dynamically switching connections on and off as needed. Indeed, excessive synchronization has been associated with pathological hypersynchrony, such as in epilepsy, whereas abnormally reduced synchronization has been linked to the functional disconnection observed in disorders such as schizophrenia [17]. These findings suggest that brain function is maintained by a dynamic balance between integration and segregation. Although the importance of this balance may appear self-evident, quantitative capture and experimental demonstrations remain challenging. Applied to visual hemifield integration, this framework suggests that, under conditions of low integration demands, information processing may be optimized through the segregation of interhemispheric connections.

In this study, we investigated the reconfiguration of EEG phase synchronization networks using a multiple object tracking (MOT) task [9], which allows for the systematic manipulation of integration and segregation demands. In this task, when stimuli moved across the left and right visual fields, the demand for interhemispheric integration increased. In contrast, when stimuli remained within a single visual hemifield, processing was confined to one hemisphere, reflecting a relatively segregated state. By leveraging this task structure, we aimed to identify EEG phase synchronization networks that are reconfigured according to integration and segregation demands. To this end, we adopted a data-driven analytical approach that did not rely on prior assumptions regarding frequency bands or electrode pairs, and analyzed phase synchronization across all electrode pairs and frequency bins. Furthermore, we examined how interindividual differences in network strength—quantified from phase synchronization—relate to behavioral performance, to obtain initial insights into individual variability in large-scale functional network dynamics under different demands for integration and segregation.

## Methods

### Participants

We recorded behavioral performance and EEG during the task to evaluate functional networks associated with interhemispheric visual integration. Sample size was estimated using G*Power [18] based on an independent preliminary dataset from 11 healthy right-handed adults (mean age ± SD: 29.6 ± 9.19 years, range: 20.0–53.0 years, 7 males). The effect size was estimated as 0.63 based on gamma-band (46–70 Hz) PSI between PO7 and PO8. This indicated that at least 36 participants were required to detect a condition difference with α = 0.05 using a two-tailed paired *t*-test.

A total of 40 healthy right-handed adults participated (25.3 ± 5.50 years, range: 18.0–38.0, 13 males). The PO7–PO8 gamma-band comparison in the full dataset was not significant (*t*(39) = -1.186, n.s.). Therefore, we adopted a whole-brain, data-driven approach using PSI across all electrode pairs and frequencies (3–57 Hz), rather than a predefined electrode pair.

All participants were naïve to the task, had normal or corrected-to-normal vision, and were right-handed according to the Japanese FLANDERS questionnaire [19,20].

The study followed the Declaration of Helsinki and was approved by the ethics committee of the National Institutes of Natural Sciences. All participants provided written informed consent. Data and analysis scripts are publicly available (https://osf.io/zmvwd).

### Multiple Object Tracking task

Two conditions were established by restricting object movement using an internal boundary (Figure 1). In the Within condition, the object movement was confined within each visual hemifield, reducing the need for interhemispheric visual integration. In contrast, in the Between condition, objects moved between the left and right visual hemifields, increasing integration cost. At the beginning of each trial, the participants memorized the target objects and were presented with a cue specifying the movement condition. During tracking, the participants tracked four target objects among a total of eight identical white objects while maintaining fixation on the central cross. At the end of each trial, the participants selected four objects using a computer mouse; if uncertain, they were instructed to guess. Visual feedback was provided, with correct selections shown in green and incorrect selections shown in red. The trial sequence is illustrated in Figure 1A. The trials were divided into six blocks, each consisting of 16 trials (eight trials per task condition, with the order randomized within each block). The participants could take breaks freely between blocks and completed one practice block before the task.

**Figure 1.**
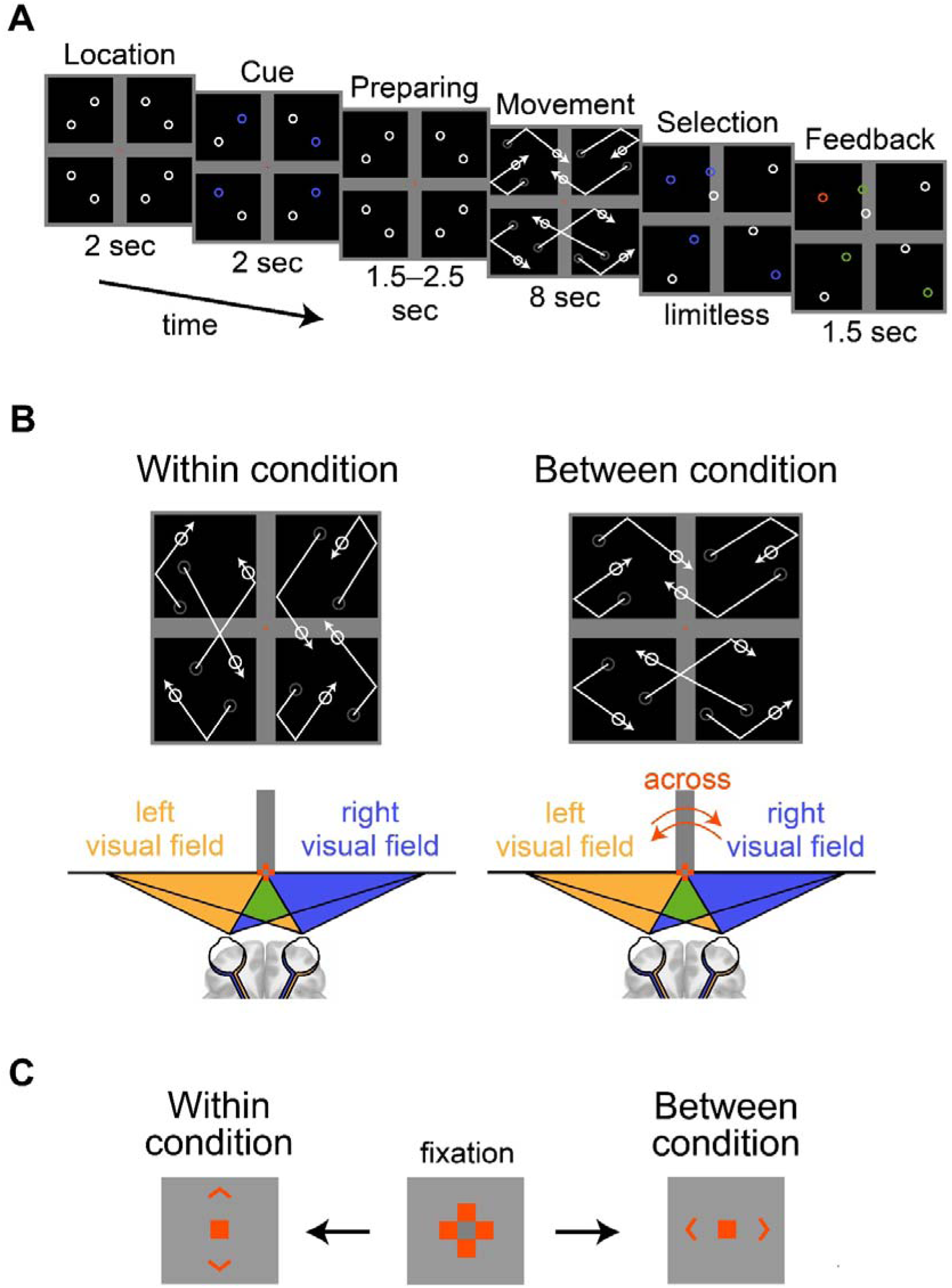
The information on the MOT task. (A) Schematic illustration of the task sequence. Participants fixed on the red cross at the center of the screen while tracking 4 targets. After the object movement ended, participants selected 4 objects they believed to be the targets using a mouse. Correct and incorrect responses were indicated in green and red. (B) The task conditions of the MOT task. In the Within condition, objects could not cross the central vertical gray boundary, whereas in the Between condition, they could not cross the horizontal internal boundary. (C) The cue difference between conditions. In the Within condition, arrows are presented in the vertical direction, whereas in the Between condition, they are presented in the horizontal direction.

### Experimental Environment

The experimental room was electromagnetically shielded and darkened to minimize visual effects other than those projected onto the PC monitor. Participants were seated 735 mm in front of a 24.5-inch LCD monitor (1920 × 1080 resolution, 100 Hz refresh rate) with their chin on a rest to maintain a consistent posture. The MOT task set-up was largely based on Bland et al. [9]. The presentation area was a square region of 24 Degrees of Visual Angle (DVA). The internal horizontal and vertical bars were 2.5 DVA in width. The fixation cross was composed of a central grey square (0.2 DVA in width) surrounded by four red squares of the same size. This cross is superior for maintaining stable fixation [21]. The circular objects were 1 DVA in diameter with 0.2 DVA white outline and moved at 10 DVA/s, as in previous studies [9].

All objects moved linearly, reflected at square boundaries or internal barriers, and passed through each other without collisions. To facilitate transitions between the quadrants, the initial movement direction was constrained within predefined angular ranges. Specifically, in the Between condition, the angle between its movement vector and the horizontal axis was restricted to ±30–45° and ±135–150°, whereas in the Within condition, it was restricted to ±45–60° and ±120–135°. The angles were randomly determined on each trial. The objects were unable to pass through the horizontal bar in the Between condition or the vertical bar in the Within condition (Figure 1B), and their movement was adjusted to prevent overlap at the end of tracking. The stimulus presentation and cursor recording were controlled using Psychtoolbox-3 [22–24].

Additionally, the signal from a photo sensor (Brain Products GmbH) was simultaneously recorded to align the stimulus timing across trials.

### Eye Tracking

Eye movements were recorded using the EyeLink 1000 PLUS (SR Research) at 2,000 Hz. The gaze coordinates were recorded during the 8-second tracking period (see Figure 1A). Trials in which the gaze deviated from beyond the vertical bar (±1.25 DVA) for more than 4 seconds were excluded from EEG analysis.

### Electroencephalography

A multichannel EEG amplifier system (ActiCHamp, Brain Products GmbH) was used to record neural activity during the task. EEG signals were recorded from 63 scalp electrodes arranged according to the international 10/10 system and were embedded in a custom-fit elastic cap (actiCAP, Brain Products GmbH), with the ground electrode at AFz. All signals were referenced the right earlobe, and the left earlobe electrode was recorded for offline re-referencing to the averaged earlobe signal. Skin/electrode impedance was maintained below 10 kΩ using an electrolyte gel (SuperVisk, EasyCap). EEG signals were sampled at 5,000 Hz and recorded using BrainVision Recorder (Brain Products GmbH). Additionally, EEG during a 3-minute eyes-open resting state were also acquired.

### Data analysis

Behavioral data were analyzed using custom MATLAB scripts (2021b, MathWorks). EEG data were analyzed using EEGLAB2021.1[25] and custom MATLAB scripts.

### Behavioral data

In each trial, the proportion of correctly identified targets (out of four) was defined as the correct rate. The trial-averaged correct rates were calculated for each condition, yielding one value per participant and condition. Differences between the conditions were tested using a one-sided paired t-test (α = 0.05).

To assess individual differences, a split-half reliability was evaluated by correlating the odd- and even-numbered trials within each condition. Intraclass correlation coefficients (ICC(2,1)) were computed to estimate the absolute agreement between single measurements under a two-way random effects model, where both participants and measurement occasions were considered random samples [26]. To visualize the relationship without assuming directional dependence, orthogonal regression based on principal component analysis (PCA) was used, with the first principal component defining the regression line.

To evaluate individual differences across conditions, we examined the relationship between trial-averaged accuracies in the Within and Between conditions and applied PCA to this two-dimensional behavioral space. The first principal component (PC1) captured the overall performance, whereas the second principal component (PC2) reflected the relative difference between conditions. Positive PC2 scores indicated better performance in the Between condition, relative to the Within condition, and negative scores indicated the opposite. PC2 score was used in neurobehavioral analyses to examine its association with EEG phase synchronization network strength.

### Preprocessing for EEG data

EEG signals were re-referenced to the averaged earlobes and baseline-corrected. A bandpass filter (1–70 Hz) and 60 Hz notch filter were applied. EEG data were segmented into 15.5-second epochs (4.5-s before to 3-s after the 8-s tracking period; Figure 1A). Independent component analysis (ICA) was conducted using the runica function in the EEGLAB to remove artifacts. Artifact-related components were identified using ICLabel [27]. Epochs exceeding ±100 microV were also removed as residual artifacts. Finally, current source density (CSD) transformation was applied using the spherical spline algorithm (CSD toolbox, version 1.1) [28,29] to reduce volume conduction effects.

### Phase synchronization analysis

To quantify the functional networks during visual field integration, we examined EEG phase synchronization. Instantaneous phase was computed using a complex Morlet wavelet transform [32] applied to CSD transformed EEG signals. The wavelet w(*t*) was defined as a complex sine wave multiplied by a Gaussian window.

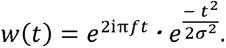

where *i* is the imaginary unit, *t* represents the time points, and *f* is the center frequency of the Gaussian window. The standard deviation of the Gaussian windowσ is defined as:

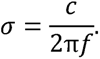

Here, *c* represents the number of cycles of the sine wave contained in the wavelet and *c* = 3 was set in this study because of higher time resolution [33]. The frequency *f* was increased from 1 to 57 Hz in 1-Hz steps. For each electrode pair, instantaneous phaseφ(*t*) was computed as:

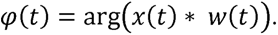

Where the EEG signal x(*t*) was convolved with w(*t*). Then, phase synchronization index (PSI) [33,34], was computed as:

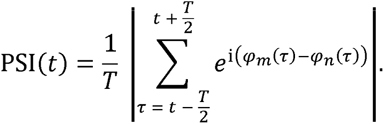

Here, *T* represents the total number of time points corresponding to the width of the time window, φ*_m_* and φ*_n_* denote the instantaneous phases at electrodes *m* and *n* at the time pointτ. The time window was slid in 20-ms steps.

### A frequency-extended network-based statistic

To enable network analysis without predefined frequency bands, we used a frequency-extended version of the network-based statistic (NBS) [36], which extends the conventional NBS framework by allowing clusters to span adjacent frequency bins. This approach enables network analysis across a continuous frequency range without requiring a priori frequency selection. EEG electrodes were defined as nodes, PSI between electrodes as links, and clusters as groups of links sharing common nodes.

A one-sided paired t-test was performed on PSI values across all links to identify condition differences. Significant links were binarized and represented in a ch × ch connectivity matrix (Figure 2A). Links sharing common nodes were grouped into clusters. This process was performed independently at each frequency. The clusters detected at adjacent frequencies were then compared, and those sharing at least one common link were merged into a single cross-frequency cluster (Figure 2B). If no shared links were present, clusters remained frequency-specific. For each detected cluster, cluster statistics were defined as the sum of the t-values of its constituent links. Statistical significance was assessed using permutation testing with 5,000 surrogate datasets, generated by shuffling task condition labels within each participant. For each permutation, only the maximum cluster statistics were obtained to construct the null distribution, and cluster statistics exceeding the top 5% threshold were considered significant.

**Figure 2.**
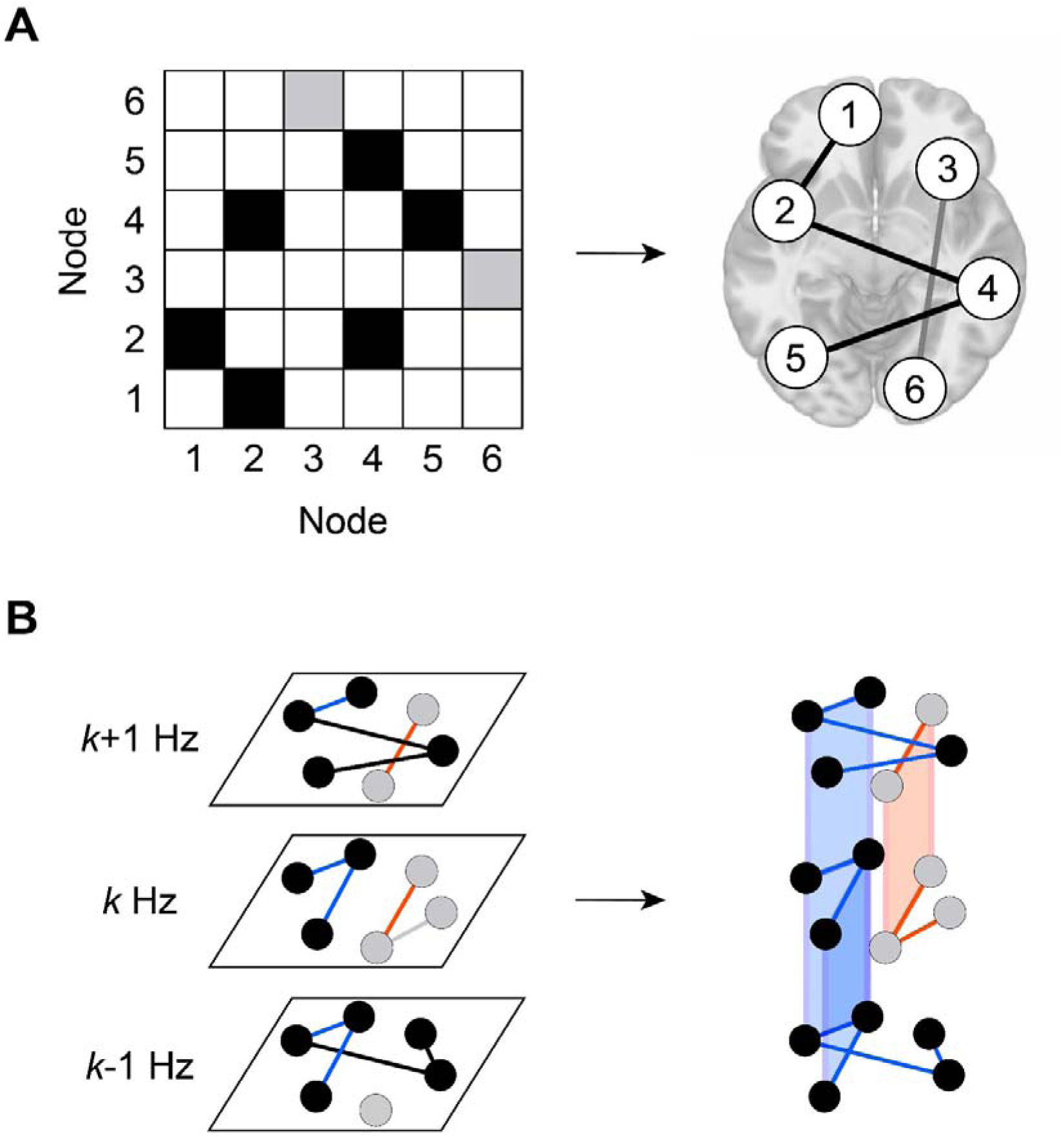
Clustering process across adjacent frequencies in the frequency-extended NBS. (A) Cluster detection in a single frequency. Only links that were significant based on statistical testing were grouped as a single cluster based on shared nodes. The same process was applied to all other frequencies. (B) Detection of cross-frequency clusters. Clusters detected at adjacent frequencies were merged if they share common links. The common links are shown in the same color, and merged clusters are illustrated in red and blue.

### Cluster network analysis

The frequency-extended NBS method was applied to the real data. A one-sided paired t-test was performed on the PSI values to compare task conditions at a significance level of 2.5% across all 1,891 electrode pairs. The analyzed frequency range was 3–57 Hz, and all other parameters were maintained at default settings.

To further examine the frequency structure of the detected network, hierarchical clustering was applied to group frequencies based on the spatial distribution patterns of their associated links. Cosine distance (1 – cosine similarity) and the average linkage method were used, and a threshold of 0.8 defined data-driven frequency groups (pseudo-frequency bands).

### Neuro-behavioral relationship

The aim of this analysis was to examine whether the networks identified in previous section were related to interhemispheric visual information integration by testing the relationship with individual differences in task performance. The trial-averaged task performance (Section “Behavioral data”) and the sum of the PSI values (cluster PSI: cPSI) within the detected network were used as representative measures. The difference between the Between and Within conditions was calculated for both measures. Two participants whose cPSI values exceeded ± 3SD were excluded from further analysis.

First, the condition differences (Between — Within) were computed for both behavioral performance and cPSI, and their correlations were examined. Next, PCA was performed separately to behavioral performance and cPSI across conditions. The PC2 scores were extracted to capture relative differences between conditions, and the correlation between the PC2 scores of behavioral performance and cPSI were analyzed. The same analysis was conducted in subgroups classified by the relative behavioral performance: participants with higher performance in the Within condition (Group W) and those with higher performance in the Between condition (Group B). Associations between behavioral and neural measures were assessed using Spearman’s rank correlation coefficients. Statistical significance was evaluated using two-tailed tests, with Bonferroni correction applied where appropriate.

### Results Behavioral results

We analyzed the differences in trial-averaged behavioral performance between the Within and Between conditions. As shown by the blue line in Figure 3A, the mean performance in the Between condition was significantly lower than that in the Within condition (*t*(37) = 1.76, *p* < 0.05), replicating a prior study (15). Despite this group-level difference, 15 out of 38 participants showed a higher correct rate in the Between condition, as indicated by the red lines in Figure 3A.

**Figure 3.**
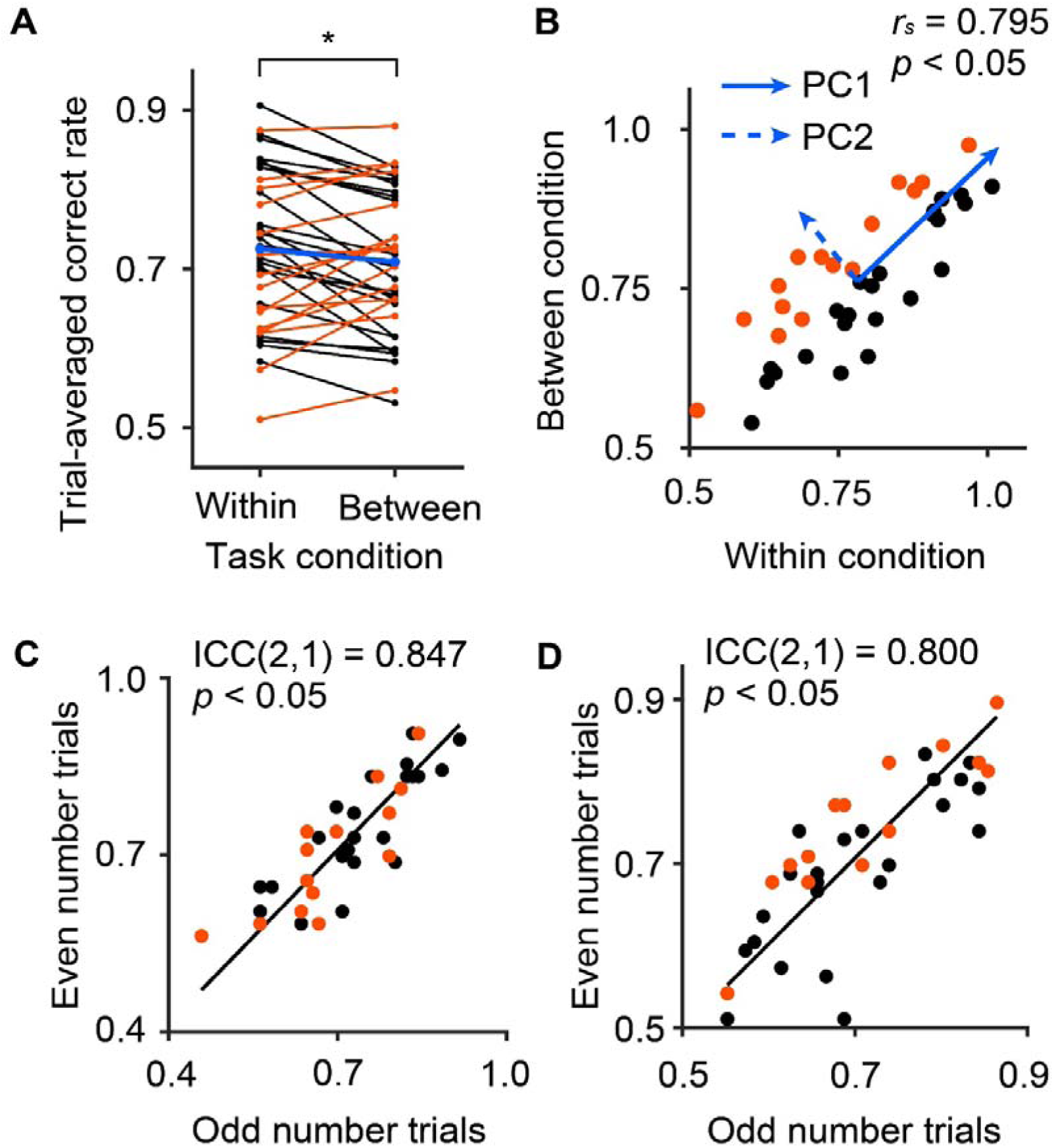
Trial-averaged behavioral performance and reliability in the MOT task. (A) Trial-averaged behavioral performance for each participant in the task conditions. The lines connected the results of same participant. The group-averages are indicated in blue. (B) Relationship between behavioral performance in the Within and Between conditions. PC1 (solid) and PC2 (dashed) indicate the principal component axes. (C)(D) Odd–even split-half reliability based on odd- and even-numbered trials in the (C) Within and (D) Between conditions.

We also examined the relationship between behavioral performance in the Within and Between conditions. As shown in Figure 3B, each participant’s correct rate under the two conditions was plotted in two-dimensional space. A significant positive Spearman’s rank correlation was observed between the two conditions (*r_s_* = 0.795, *p* < 0.05), indicating that individuals who performed well in one condition tended to perform well in the other condition. To further characterize the structure of individual differences in this behavioral distribution, we applied PCA. The resulting principal component vectors (PC1 and PC2) are overlaid in the figure to illustrate the primary axes of variance among participants.

To assess the odd–even split-half reliability of interindividual differences in behavioral performance within each task condition, we conducted an odd–even split-half reliability analysis based on the consistency between odd-end and even-numbered trials. Because both variables are subject to measurement errors and no directional dependency was assumed, the regression line shown in Figure 3C and 3D was defined as the first principal component axis (orthogonal regression) of the two-dimensional data. In the Within condition, the orthogonal regression line comparing odd- and even-numbered trials was Y = 0.982 X + 0.0209. The intraclass correlation coefficient (ICC(2,1)) indicated a significant reliability (ICC(2,1) = 0.847, *p* < 0.05), as shown in Figure 3C. In the Between condition, the orthogonal regression line was Y = 1.05X – 0.0260, and the ICC(2,1) also indicated a significant reliability (ICC(2,1) = 0.800, *p* < 0.05), as shown in Figure 3D. These p -values were derived from the standard F-test used in the ICC computation.

Participants with higher performance in the Within condition are shown in black, and those with higher performance in the Between condition are shown in red. Participants whose trial-averaged correct rate was higher in the Within condition are shown in black, while those with higher correct rate in the Between condition are shown in red in (A), (B), (C), (D)

### The results of the frequency-extended NBS

We applied a frequency-inclusive data-driven extension of the NBS to identify functional networks showing condition-related differences in phase synchronization. PSI values were computed for each electrode pair and frequency (3–57 Hz), and statistical comparisons Within and Between conditions were performed for each frequency–pair combination. Suprathreshold links were grouped into clusters based on their spatial and frequency adjacencies. A single large cluster spanning a broad frequency range emerged, indicating consistently greater PSI in the Between condition across multiple frequencies.

To characterize the spatial-frequency structure of the detected network, we examined the similarity of the spatial link patterns across frequencies. Hierarchical clustering using cosine distance grouped frequencies into seven groups (Figure 4A). The groups did not align with the conventional frequency bands but formed continuous range of adjacent frequencies. We refer to them as "pseudo-frequency bands"

**Figure 4.**
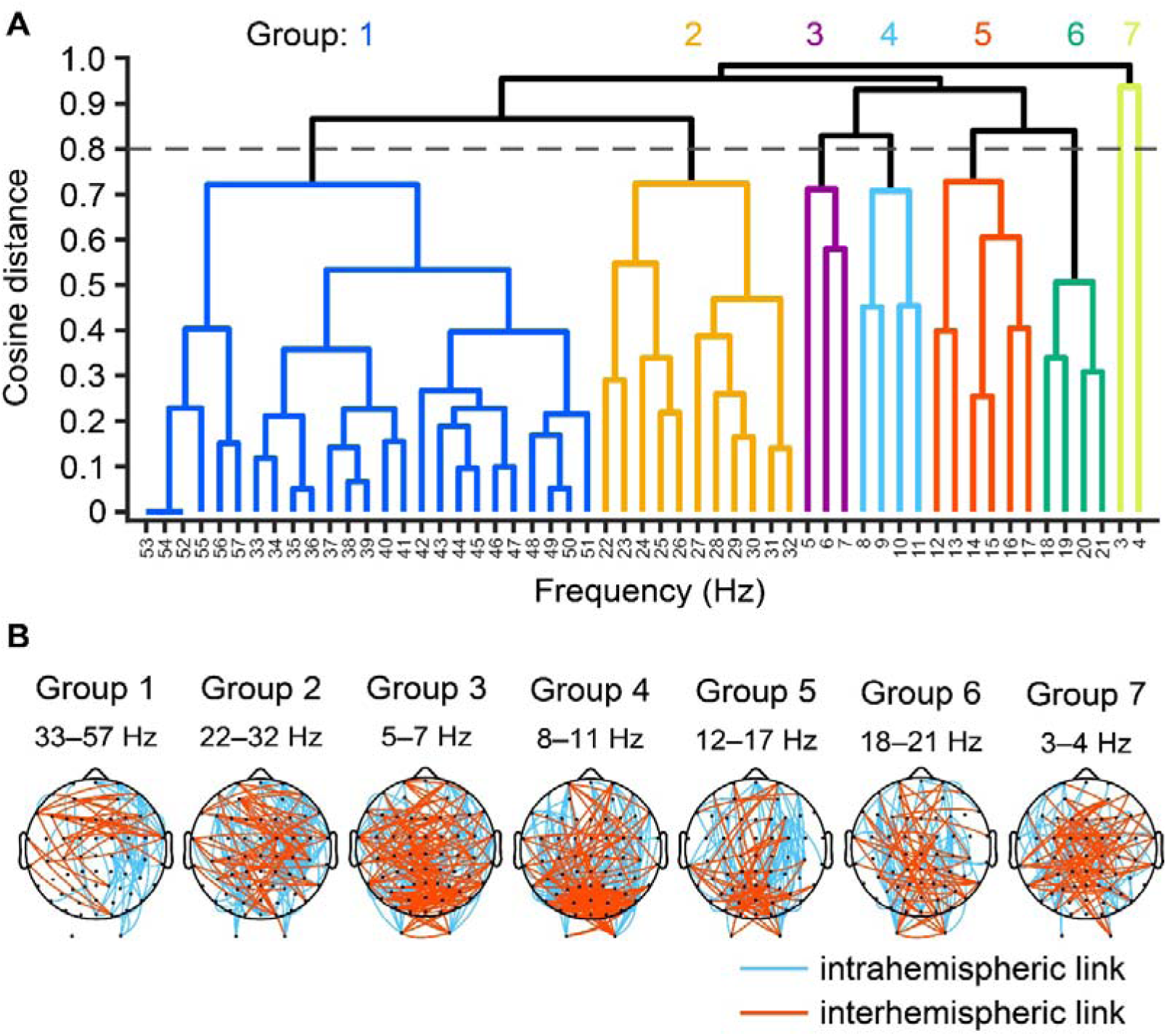
Multi-frequency architecture of the frequency-extended NBS-detected synchronization network. (A) Hierarchical clustering of frequencies based on the similar network structures, yielding seven data-driven frequency groups. (B) Spatial distribution of network links for each frequency group. Red and light blue lines indicate interhemispheric and intrahemispheric connections, respectively.

To visualize the spatial structure, network links were plotted for each pseudo-frequency band (Figure 4B). Groups 2 (22–32 Hz), 6 (18–21 Hz), and 7 (3–4 Hz) showed broadly distributed links, whereas Group 1 (33–57 Hz) exhibited prominent interhemispheric synchronization localized primarily in the frontal regions. Groups 3 (5–7 Hz), 4 (8–11 Hz), and 5 (12–17 Hz) displayed dense interhemispheric connectivity in the occipital areas. All pseudo-frequency bands included long-range fronto-occipital intrahemispheric links, with Groups 1 and 5 exhibiting right-lateralized patterns.

### Neuro-behavioral relationship in the detected network

To interpret the functional relevance of the detected network, we investigated whether individual variability in network strength was associated with behavioral performance. Specifically, we focused on condition-related changes (Between – Within) in both neural and behavioral measures to test whether greater neural modulation corresponded to greater behavioral modulation.

Network strength was quantified as the sum of the PSI values across all links in the detected network, yielding a condition-specific cPSI score (Figure 5A). Relationships between Within and Between conditions were visualized using scatter plots with PCA (Figure 5B). In this plot, PC1 captures the overall degree of cPSI across conditions, whereas PC2 represents the relative difference between conditions, indicating the extent to which cPSI is greater in the Between condition than in the Within condition. We then computed the condition differences (Between – Within) for both cPSI and behavioral performance and examined their Spearman’s rank correlation (Figure 5C). This difference-based approach treats the magnitude of change as an individual marker of sensitivity to interhemispheric integration costs. No significant correlation was observed (*r_s_*= –0.0355, n.s.).

**Figure 5.**
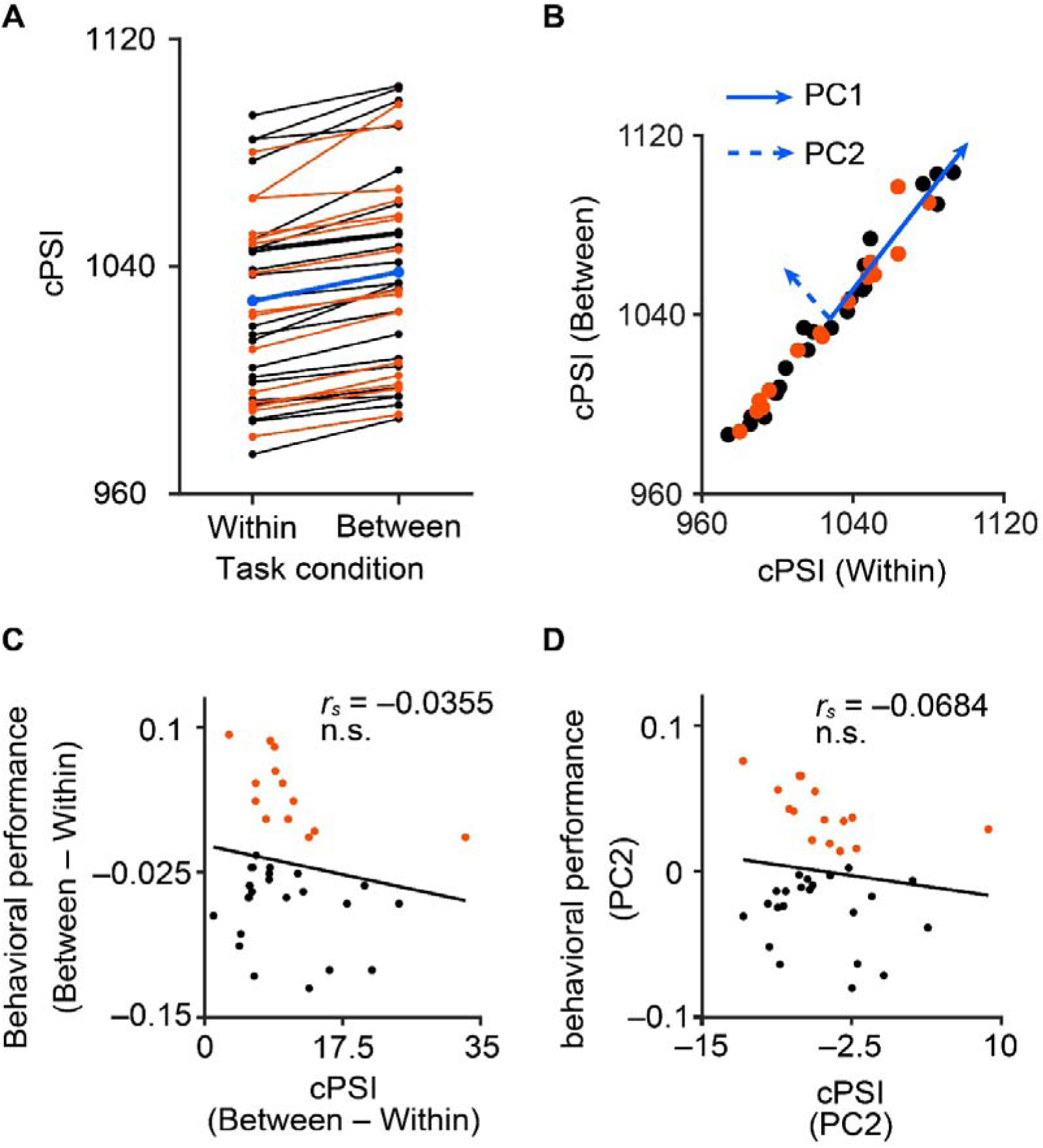
Intra- and interindividual variability in phase synchronization network strength and its association with behavioral performance. (A) The cPSI in the Within and Between conditions. The results of the same participant connected by a line. (B) Relationship between cPSI in the Within and Between conditions with PC1 (solid) and PC2 (dashed). (C) Correlation between the difference in behavioral performance and cPSI (Between – Within). (D) Correlation between PC2 scores of behavioral performance and cPSI. In (C) and (D), the solid line represents the orthogonal regression (see Methods). In all figures, participants whose trial-averaged correct rate was higher in the Within condition are shown in black, while others are shown in red.

Additionally, we extracted PC2 scores from the PCA of both behavioral performance (Figure 3B) and cPSI and examined their correlations (Figure 5D). The PC2 score represents the primary axis of condition-related modulations, capturing the relative difference in network strength between the conditions: Participants with positive PC2 scores exhibited stronger phase synchronization (higher cPSI) in the Between condition than in the Within condition. The correlation between the PC2 scores of behavioral performance and cPSI was not significant (*r_s_* = –0.0684, n.s.).

Next, to examine consistency with prior findings, we conducted an additional analysis using only the interhemispheric links within the frequency-extended NBS-identified network. Because visual hemifield integration depends on interhemispheric connections [8,9], fluctuations in EEG phase synchronization across these links are likely related to task performance.

When all participants were considered, a significant positive Spearman’s rank correlation was observed between behavioral and cPSI_IntHem difference scores (Between – Within) (*r_s_* = 0.420, Bonferroni-corrected *p* < 0.05; Figure 6A). Similarly, a significant positive correlation was found for PC2 scores (*r_s_* = 0.425, Bonferroni-corrected *p* < 0.05; Figure 6B). Similarly, to test whether these associations were specific to interhemispheric connections, we further conducted the analysis using only intrahemispheric links. In contrast, neither the difference scores (Between – Within) (*r_s_* = 0.115, n.s.; Figure 6C) nor PC2 scores (*r_s_* = 0.112, n.s.; Figure 6D) were significantly associated with intrahemispheric cPSI.

**Figure 6.**
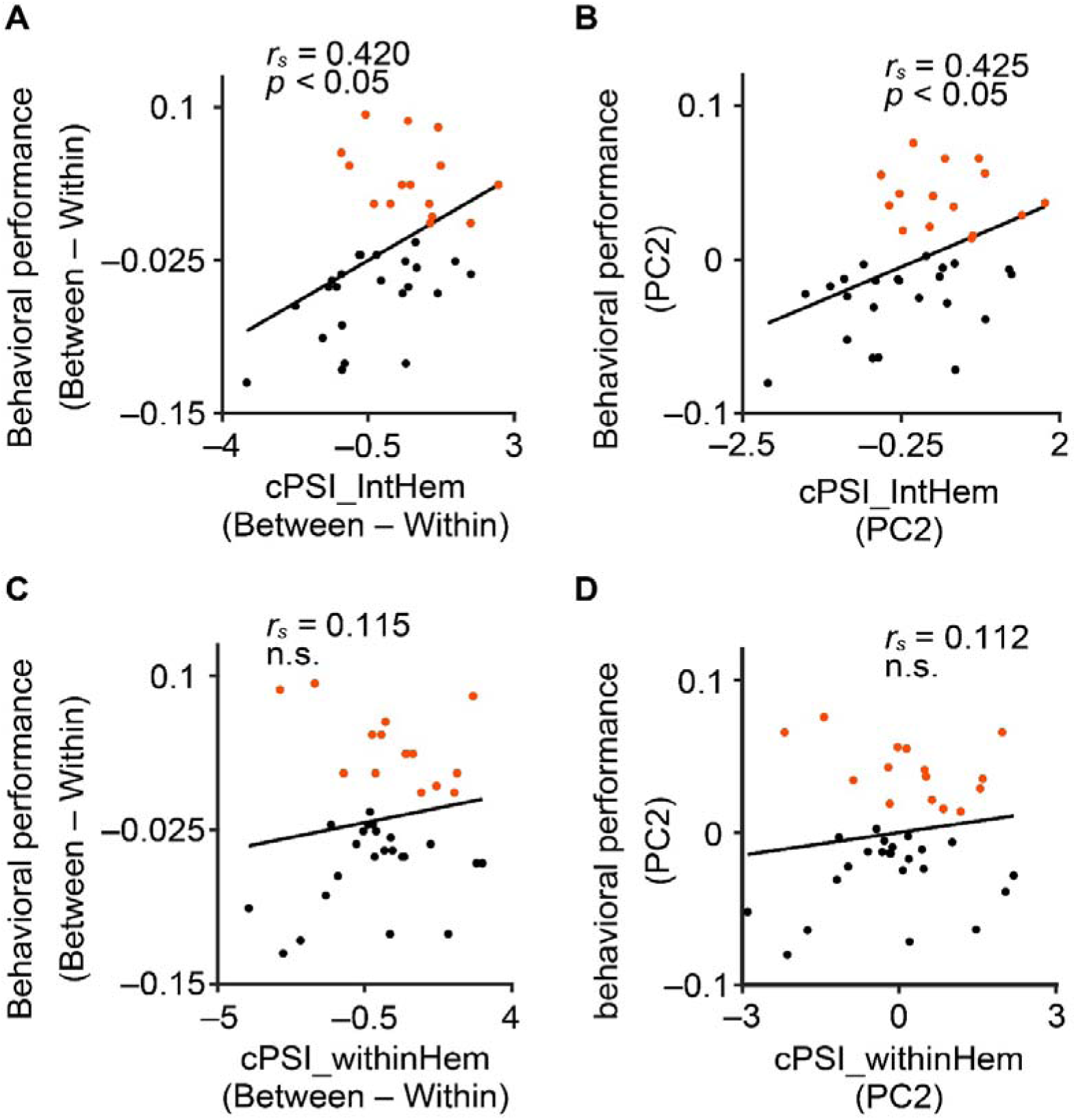
Relationship between behavioral performance and EEG phase synchronization based on interhemispheric and intrahemispheric connections within the frequency-extended NBS-identified network. (A–D) The same analysis as in Figure 5C and 5D, but using only the (A–B) interhemispheric links / (C–D) intrahemispheric links contained within the significant cluster identified by the frequency-extended NBS. (A, C) Correlation between condition differences (Between − Within). (B, D) Correlation between PC2 scores. Solid lines indicate orthogonal regression (see Methods); significance was assessed using Spearman’s correlation. Black and red points denote participants with higher performance in the Within and Between conditions, respectively.

To examine whether the relationship between behavioral performance and network strength varied across participants, we conducted a subgroup analysis based on relative performance. Participants were divided into group W and group B.

For each group, the condition differences in cPSI_IntHem and behavioral performance were examined and correlated. In Group W, a significant positive correlation was observed (*r_s_* = 0.448, Bonferroni-corrected *p* < 0.05; Figure 7A), whereas no significant correlation was observed in Group B (*r_s_* = –0.405, n.s.; Figure 7B). For PC2 scores, neither group showed a significant relationship (Group W: *r_s_* = 0.305, n.s.; Group B: *r_s_* = –0.0964, n.s.; Supplemental Figure 1A–B). To examine whether similar subgroup-dependent patterns were present for interhemispheric links, the same analyses were conducted using cPSI_withinHem. In contrast to interhemispheric results, no significant correlations were observed in either group for difference scores (Group W: *r_s_* = 0.0654, n.s.; Group B: *r_s_* = –0.457, n.s.; Figure 7C–D), or the PC2 scores (Group W: *r_s_* = 0.0356, n.s.; Group B: *r_s_* = –0.371, n.s.; Supplemental Figure 1C–D).

**Figure 7.**
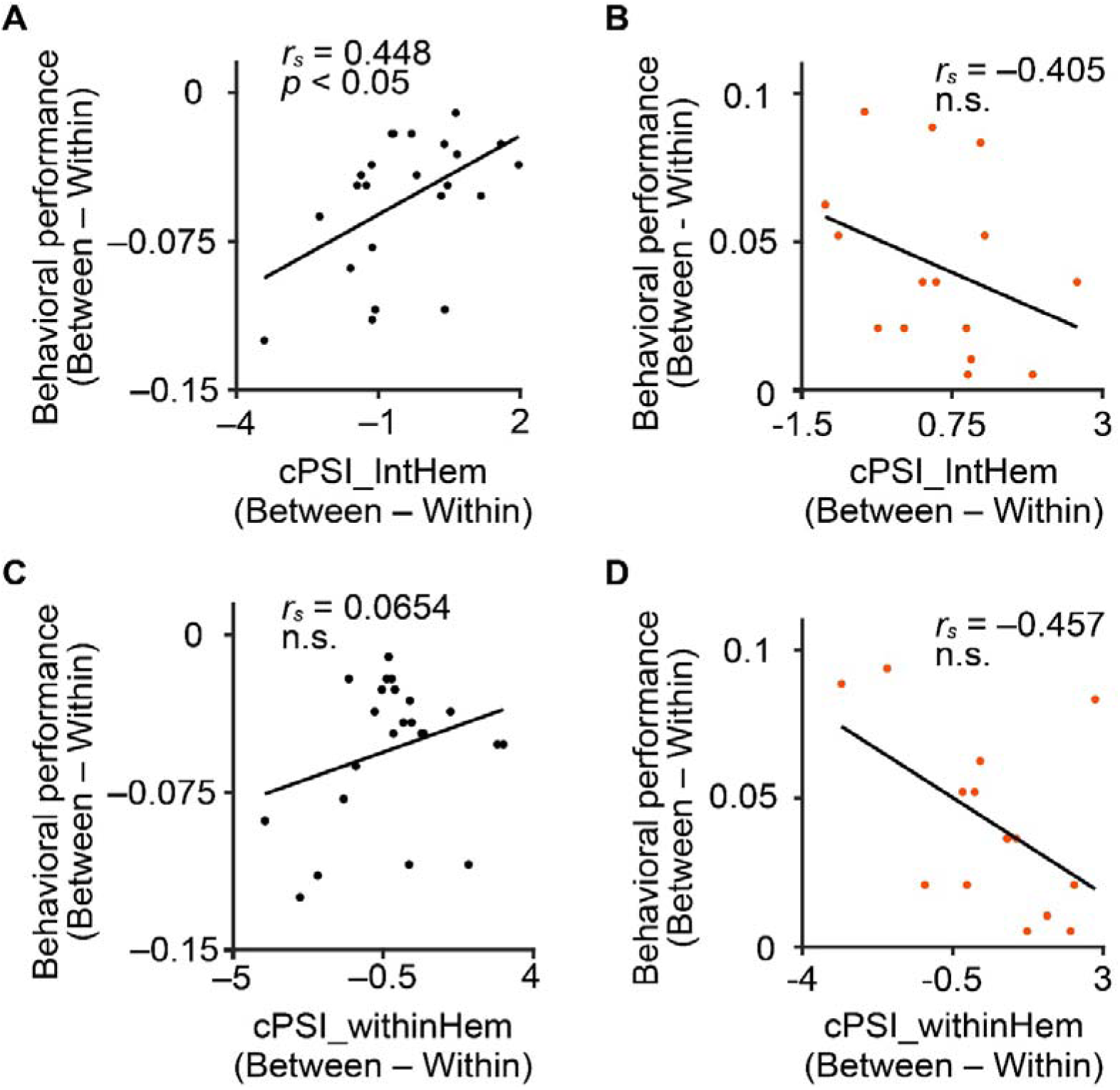
Subgroup-specific relationships between behavioral performance and integration–segregation contrast in EEG phase synchronization. (A–D) The relationship between cPSI and behavioral performance during the Between condition, based on the difference from the Within condition. Only the (A,B) interhemispheric / (C,D) intrahemispheric links within the frequency-extended NBS-identified clusters were used. (A,C) show the results for group W, and (B,D) show the results for group B. In all panels, the black dots represent the participants in group W, and the red dots represent those in group B. The solid line represents the orthogonal regression line shown in Figure 6.

## Discussion

### Behavioral cost and individual differences associated with visual hemifield integration

Participants showed lower task performance in the Between condition than in the Within condition, consistent with previous studies demonstrating increased processing costs when tracking objects across visual hemifields [37,38]. Because the two conditions were matched in spatial distribution and task structure except for midline crossing, this difference is likely attributable to increased demands on interhemispheric integration [39,40]. This behavioral difference provides an important foundation for examining the relationship between behavioral performance and subsequent neural activity.

However, this tendency was not uniform across participants. A substantial number of individuals (15/40) showed numerically higher performance in the Between condition, highlighting stable interindividual variability rather than measurement noise, as supported by high Odd–even split-half reliability. Importantly, such variability cannot be explained solely by a single axis of having “stronger or weaker integration capacity.” Instead, it is more consistent with theoretical frameworks emphasizing the balance between integration and segregation [14–16]. Thus, the individual differences in performance likely reflect variability not only in integration ability but also in the capacity for functional segregation.

### Large-scale EEG phase synchronization network during visual field integration

The frequency-extended NBS analysis revealed a large-scale network exhibiting significantly stronger EEG phase synchronization only in the Between condition. Notably, this network was characterized by a prominent involvement of interhemispheric connections, consistent with the increased demand for integrating visual information in the Between condition. This finding supports the interpretation that the detected network reflects a functional substrate underlying visual hemifield integration. In addition, because the frequency-extended NBS defines clusters as groups of links sharing at least one node, the detected network also included several intrahemispheric connections. This suggests their contribution to visual hemifield integration [7].

Importantly, no significant clusters showing the opposite pattern—that is, stronger EEG phase synchronization in the Within condition—were detected. Together, these results suggest that visual hemifield integration is supported by the dynamic recruitment of large-scale EEG phase synchronization networks spanning multiple cortical regions. This pattern further implies that a baseline network common to both conditions may exist, with minimal intrahemispheric modulation in the Within condition, whereas both inter- and intrahemispheric synchronization increased in the Between condition. This view aligns with the notion that the balance between integration and segregation dynamically shifts depending on cognitive state or task demands [14–16]. Therefore, large-scale EEG phase synchronization is also suggested to provide important clues for understanding the process by which the brain integrates and segregates visual information across hemispheres.

### Neuro-behavioral relationship and individual differences

We examined whether individual differences in network strength were associated with behavioral performance. As a result, a positive correlation was observed between task performance and cPSI_IntHem, which was calculated using only the interhemispheric links (Figure 6A, B). Furthermore, subgroup analyses revealed a positive correlation in Group W (Figure 7A) and a trend toward a negative correlation in Group B (Figure 7B). These results suggest that higher phase synchronization is not necessarily advantageous. Rather, the ability to flexibly increase synchronization under heightened integration demands may contribute to better performance. More specifically, Group W appeared to represent participants who could effectively segregate their networks during the Within condition and appropriately enhance synchronization during the Between condition, leading to improved performance. In contrast, Group B participants may have been unable to segregate their networks sufficiently during the Within condition and maintain a high level of synchronization, such that further increases during the Between condition became excessive and interfered with efficient integration. These findings are consistent with theoretical perspectives emphasizing that the dynamic balance between integration and segregation constitutes a fundamental principle supporting task execution [14–16,41], as well as with the importance of flexibly reconfiguring large-scale connectivity structures [42–44], providing novel experimental evidence for these frameworks.

### Functional interpretation of frequency-specific components

To further characterize the organization of the detected network, hierarchical clustering was applied to the frequency dimension of the EEG phase-synchronized links identified using the frequency-extended NBS. This analysis revealed contiguous, data-driven frequency groupings, referred to here as pseudo-frequency bands, which, although not aligned with canonical definitions, exhibited distinct spatial patterns (Figure 4A–B).

Because the frequency-extended NBS imposes no constraint on spectral adjacency, the emergence of these clusters suggests that the spatial structure of EEG phase synchronization varies gradually across frequencies, indicating an intrinsic spectral hierarchy in interhemispheric coupling. These results support analyzing EEG phase synchronization in a frequency-inclusive, data-driven manner rather than relying on predefined bands. The spatial configurations associated with each pseudo-frequency band reflected both local and long-range interactions across cortical regions. For instance, lower-frequency clusters (e.g., Groups 3 and 7) showed widespread interhemispheric connectivity, whereas higher-frequency clusters, such as Group 1, exhibited a more localized frontal dominance. This aligns with the notion that large-scale EEG phase synchronization is temporally multiplexed across multiple frequency channels [2]. Consistent with theoretical models, different oscillatory frequencies support distinct functional roles, with low-frequency oscillations (e.g., delta and theta) associated with long-range coordination and high-frequency oscillations (e.g., gamma) linked to local processing and temporal precision [10,45,46]. The widespread interhemispheric synchronization in Groups 3 (5–7 Hz) and 7 (3–4 Hz) supports their role in global integration, while posterior alpha activity in Group 4 (8–11 Hz) aligns with visual cortical gating functions.

These observations refine and extend prior findings. For example, Bland et al. (2020) reported gamma-band coherence between bilateral occipitotemporal regions during a similar MOT task [9]. Although some pseudo-frequency clusters overlapped this pattern, particularly in posterior regions, we also observed robust interhemispheric synchronization in the theta range, which was not reported by Bland et al. This theta activity may reflect increased working memory or control demands during visual hemifield integration [47,48], whereas frontal-dominant gamma synchronization in Group 1 may reflect executive monitoring and attentional regulation [49]. Overall, the spatial and spectral structures in Figure 4 indicate that cross-hemifield visual integration recruits frequency-differentiated phase synchronization networks, highlighting the value of data-driven, frequency-inclusive approaches for capturing dynamic coordination across large-scale brain networks.

### Methodological implications of frequency-spanning network detection using the frequency-extended NBS

A key methodological feature of this study lies in the application of the frequency-extended NBS, designed to detect functional connectivity clusters spanning adjacent frequencies. Unlike conventional NBS approaches [36], which require separate analyses for each frequency band, the frequency-extended NBS enables the identification of statistically significant networks across a continuous spectral range without strong a priori assumptions about frequency relevance. This is particularly valuable in cognitive paradigms such as interhemispheric visual integration, where multiple frequency bands may jointly support large-scale coordination [10]. Restricting analyses to predefined bands risks overlooking cross-frequency interactions and underestimating distributed neural dynamics. By aggregating connections that share spatial patterns across neighboring frequencies, the frequency-extended NBS maintains the core cluster-based permutation inference of NBS while improving sensitivity to complex network structures. In the present study, the frequency-extended NBS identified a single, large-scale network spanning 3–57 Hz (Figure 4), which would likely appear fragmented under frequency-specific analyses. This multifrequency detection revealed a coherent network associated with visual hemifield integration.

More broadly, the frequency-extended NBS provides a flexible and data-driven framework for the exploratory identification of multifrequency networks when spectral signatures are unknown or heterogeneous. Although this method assumes some continuity across frequencies, this trade-off is acceptable when mapping distributed network dynamics. Future extensions may integrate source-level connectivity or time-resolved clustering to enhance anatomical or temporal resolution. Nevertheless, the present implementation demonstrates that frequency-inclusive network statistics are useful for investigating large-scale functional organization.

Although the frequency-extended NBS effectively detects frequency-spanning connectivity patterns, it does not capture the temporal dynamics of network emergence. Therefore, the identified cluster reflects an overall trial-averaged effect rather than a transient synchronization change. Future studies using time-resolved or sliding-window analyses could clarify how interhemispheric synchronization evolves during visual field integration.

### Conclusions

Using a frequency-inclusive network analysis, this study showed that interhemispheric visual integration is supported by large-scale EEG phase synchronization networks spanning multiple frequency bands. Individual differences in behavioral performance were associated with how flexibly these networks were modulated, suggesting that efficient integration depends on appropriately tuned rather than maximally enhanced synchronization. These findings underscore the value of data-driven frequency-spanning approaches for studying large-scale functional networks in the human brain.

## Acknowledgments

This work was supported by JSPS KAKENHI (Grant Number JP19H04024), the JST Moonshot R&D Program (JPMJMS2292), and the NIPS Encouraging Grant for Graduate Students.

## Author Contributions

Makoto Hagihara: Conceptualization, Data curation, formal analysis, Investigation, Methodology, Software, Validation, Visualization, Writing – original draft, writing – review, and editing.

Kazumasa Uehara: Conceptualization, Supervision, Writing – review & editing Yuka Okazaki: Software, Writing – review & editing Keiichi Kitajo: Conceptualization, Methodology, Funding acquisition, project administration, Resources, Supervision, Writing – review, and editing

---

**Supplemental Figure 1.**
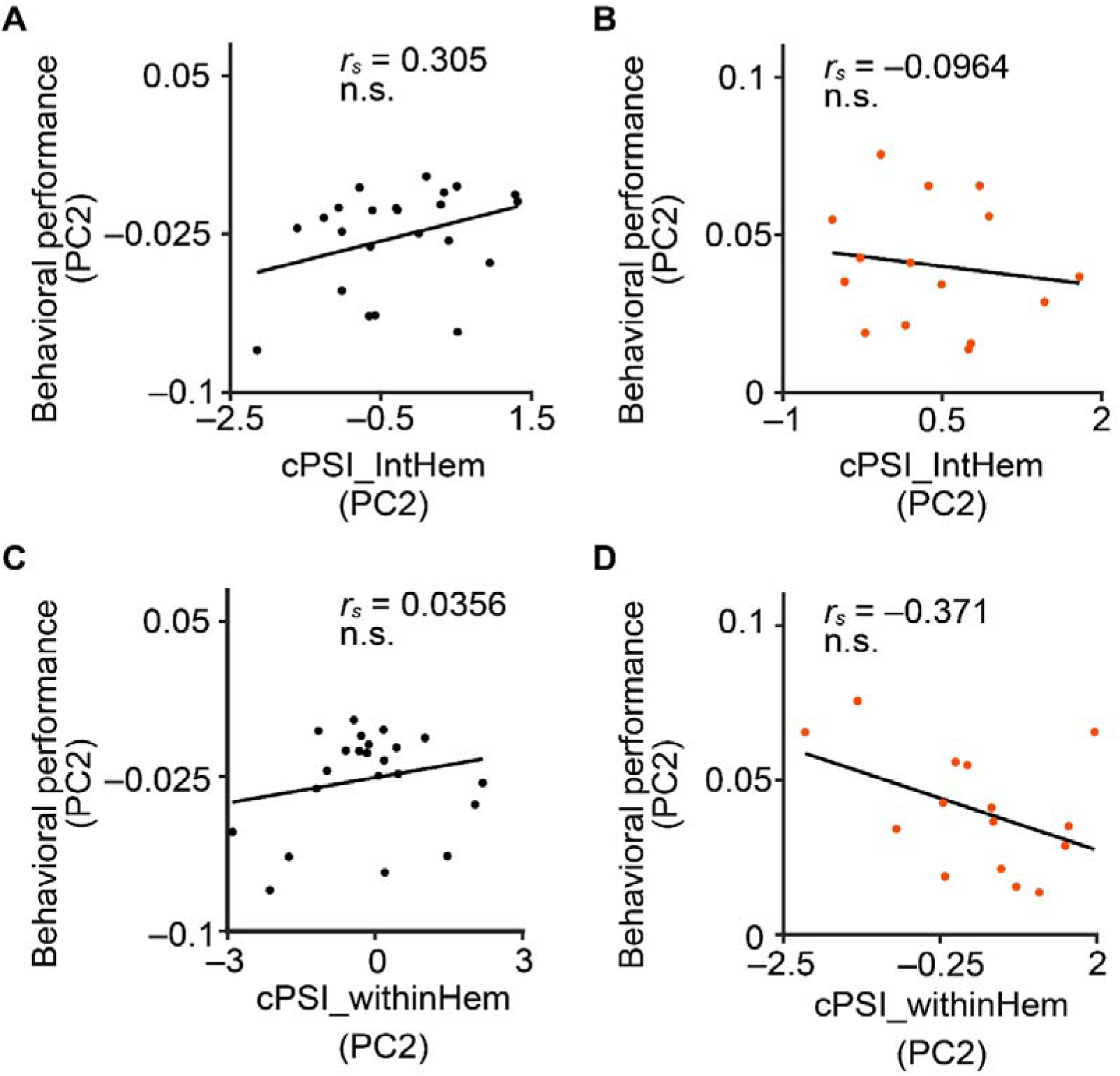
Relationship between PC2 scores of the behavioral performance and EEG phase synchronization based on the connections identified by the frequency-extended NBS during visual hemifield integration in each subgroup. (A–D) The relationship between PC2 scores of the cPSI and behavioral performance. Only the (A,B) interhemispheric / (C,D) intrahemispheric links within the frequency-extended NBS-identified cluster was used. (A,C) shows the result of group W, and (B,D) shows the results of group B. In all panels, black dots represent participants in group W, and red dots represent those in group B.

